# Single cell RNA-seq in Drosophila testis reveals evolutionary trajectory of sex chromosome regulation

**DOI:** 10.1101/2022.12.07.519494

**Authors:** Kevin H-C Wei, Kamalakar Chatla, Doris Bachtrog

**Affiliations:** Department of Integrative Biology, University of California Berkeley, Berkeley, CA 94720, USA

## Abstract

Although sex chromosomes have evolved from autosomes, they often have unusual regulatory regimes that are sex- and cell-type-specific such as dosage compensation (DC) and meiotic sex chromosome inactivation (MSCI). The molecular mechanisms and evolutionary forces driving these unique transcriptional programs are critical for genome evolution but have been, in the case of MSCI in Drosophila, subject to continuous debate. Here, we take advantage of the younger sex chromosomes in *D. miranda* (XR and the neo-X) to infer how former autosomes acquire sex-chromosome specific regulatory programs using single-cell and bulk RNA sequencing and ribosome profiling, in a comparative evolutionary context. We show that contrary to mammals and worms, the X’s are downregulated through germline progression most consistent with a loss of DC instead of MSCI, resulting in half gene dosage at the end of meiosis for all three X’s. For the young neo-X, DC is incomplete across all tissue and cell types and this dosage imbalance is rescued by contributions from Y-linked gametologs which produce transcripts that are translated to compensate both gene and protein dosage. We find an excess of previously autosomal testis genes becoming Y-specific, showing that the neo-Y and its masculinization likely resolve sexual antagonism. Multicopy neo-sex genes are predominantly expressed during meiotic stages of spermatogenesis, consistent with their amplification being driven to interfere with Mendelian segregation. Altogether, this study reveals germline regulation of evolving sex chromosomes and elucidates the consequences these unique regulatory mechanisms have on the evolution sex chromosome architecture.

## INTRODUCTION

Sex chromosomes often show unusual expression patterns relative to autosomes. Repeat-rich and gene-poor Y chromosomes are often epigenetically silenced [1–3]. X chromosomes, in response, have evolved dosage compensation in many species to equalize expression of the sex chromosomes and autosomes in both sexes [4–7]. This is accomplished via diverse mechanisms found in different taxa including down regulation of one copy of the X in female mammals and reduced expression of both copies in worms [5,8,9]. In Drosophila, dosage compensation occurs via hyper transcription of the single X in males. This is achieved via the sequence-specific recruitment of the male-specific lethal (MSL) complex to so-called chromatin entry sites (CES) on the X which induces the activating histone tail modification H4K16 acetylation (H4K16ac) at nearby transcribed genes [10–12].

Germline regulation of sex chromosomes also appears to be uniquely distinct from autosomes. In many organisms, expression on the X is highly reduced during meiosis of spermatogenesis, a phenomenon known as meiotic sex chromosome inactivation (MSCI) [13–15]. In mammals, the X is sequestered in a silencing compartment in the nuclear periphery known as the sex-body during late meiotic prophase [16–20]. Similarly, the germline of XO male worms creates a structure analogous to the sex-body, where the X is associated with the repressive histone tail modification H3K9-methylation [13,21]. Despite the apparent conservation, the evolutionary importance of MSCI remains unclear but MSCI may prevent unwanted recombination between the heterologous sex chromosomes [22], safeguard the meiotic progression of asynapsed chromosomes [23], silence the expression of a feminized X chromosome to reduce sexual antagonism [24] or inactivate sex-linked selfish genetic elements that distort sex ratio [25–27].

Interestingly, unlike other model organisms such as mice and worms, the status of MSCI in the male germline of Drosophila has been contentious [28–36]. In the apical tip of the fly testes, germline stem cells (GSCs) are supported by somatic hub and cyst cells. After differentiation, the spermatogonia (SG) undergo 4 rounds of cell divisions with incomplete cytokinesis resulting in multinucleated cysts called the spermatocyte (SC) that enter meiotic division. Late spermatocytes exit meiosis forming haploid spermatids (ST) that remain connected until spermiogenesis when sperm maturation, individualization, and elongation occur to form mature sperm bundles. X-linked expression across the germline appears to be reduced when assayed with qPCR of targeted genes [30,34], reporter constructs [28,34], and chromosome-wide transcriptome analyses [29–32]. However, whether this downregulation is specific to meiotic tissues or can be attributed to MSCI has been debated. Dosage compensation further confounds the interpretation of germline X-regulation in Drosophila. While MSL-dependent dosage compensation has been firmly established in somatic tissues, its presence in the testes is unclear. Cytological studies of testis have failed to detect the MSL complex or the H4K16ac modification it induces in GSC, SG, and SC [37]. However, recent genomic studies have revealed X-linked expression patterns and nuclear topologies that suggest that a distinct mechanism maintains X dosage in the premeiotic stages of spermatogenesis [20,35,38].

Gene content and gene expression evolution further complicates inferences of chromosome-wide expression patterns of sex chromosomes during spermatogenesis. Genes that are expressed in testis have been found to move off the X chromosome [39–42], while sexually antagonistic selection could result in an excess or deficiency of genes with male- or female biased expression on sex chromosomes, depending on underlying population parameters [43–46]. In particular, female-biased transmission of the X may select for an excess of female-specific genes / a deficiency of male-specific genes, and overall lower expression of the X compared to autosomes in males (“demasculinization” of the X; [43,47]). Male-beneficial genes, on the other hand, should be over-represented on the male-specific Y chromosome [43,48–50].

Because sex chromosomes typically have autosomal origins [51,52], newly sex-linked chromosomes will gradually acquire regulatory features of older sex chromosomes over evolutionary time. Due to its unusual sex chromosome architecture, *Drosophila miranda* has been a model species for understanding sex chromosome evolution (**Figure 1A**). Unlike the ancestral karyotype of Drosophila where a single chromosome arm (Muller A) is the X chromosome, *D. miranda* has two additional X chromosome arms that are younger (**Figure 1B**). A metacentric fusion linked one autosome, Muller D, with Muller A ~15MYA, causing the former to become X-linked. Muller D has evolved the stereotypical characteristics of a sex chromosome, with the non-recombining homolog almost fully degenerate, and the X-linked homolog having evolved full dosage compensation [53] (note that we refer to this chromosome arm as Muller AD, since a pericentric inversion moved some Muller A genes onto Muller D). A secondary fusion between Muller C and the Y occurred around ~1.5MYA creating a Y-linked copy of Muller C that is male exclusive (i.e. the neo-Y). Its former homolog (the neo-X), while not fused to the Muller A and D, is found as two copies in females and one copy in males. The neo-X and neo-Y chromosomes have been evolving characteristic traits like X dosage compensation [2,54] and Y degeneration [55–58], and are at an intermediate stage in the transition from an ordinary autosome to a pair of differentiated sex chromosomes [57,58]. The acquisition of dosage compensation of the neo-X occurred through co-opting insertions of a transposable element that harbors the sequence motif that is targeted by the MSL complex [53,59].

**Figure 1.**
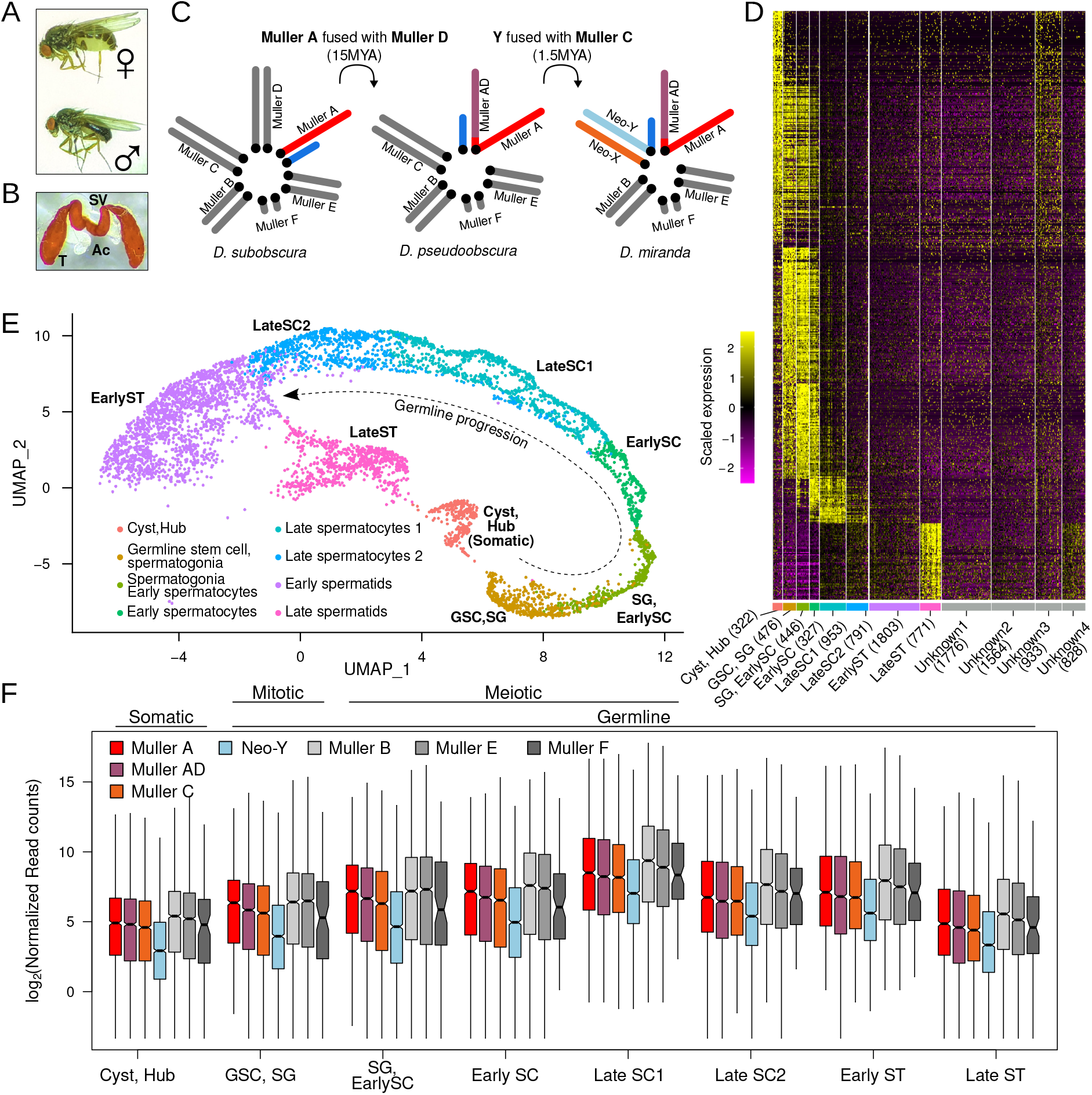
Transcriptional dynamic of D. miranda testes at single cell resolution. **A.** Picture of *D. miranda* female and male adults. **B.** Picture of a pair of testes (T) attached by the seminal vesicle (SV) to the accessory glands (Ac). **C.** Male karyotype evolution showing multiple fusions creating neo-sex chromosomes of different ages. Red, purple and orange denote the three X-chromosomes of different ages: Muller A (50MY), Muller AD (15MY), and Muller C (1.5MY), respectively. The Y chromosome arms are in shades of blue, and the autosomes are in gray. Unless otherwise stated, this color-coding of the chromosomes will be maintained throughout. **D.** Expression of genes used for clustering cells into cell types; number of cells (UMIs) in each type is in parantheses. Yellow and magenta represent high and low expression. **E.** UMAP reduced dimension projection showing cells from scRNA-seq clustered into 8 different cell types corresponding to different stages of spermatogenesis. Progression of spermatogenesis is outlined by curved arrow. For projection of all cells including unidentified clusters, see Supplementary Figure 1. **F.** Distributions of normalized gene counts from different chromosomes across cell types. Cell types are further labeled as somatic, germline, mitotic and meiotic above the plot.

Here, we employed single cell RNA-seq (scRNA-seq) on *D. miranda* testes (**Figure 1C**) to disentangle the regulatory mechanisms of sex chromosomes during spermatogenesis and to understand how young sex chromosomes evolve in response to such meiotic regulatory regimes. We find expression patterns consistent with incomplete dosage compensation of the neo-X prior to meiosis followed by loss of dosage compensation across all X’s; this suggests the loss of dosage compensation as the source of meiotic downregulation of the X. Further, comparing expression pattern of the neo-X to its autosomal ortholog in *D. pseudoobscura* using bulk tissue RNA-seq, we found that neo-X expression is reduced in all tissues, due to suboptimal dosage compensation in the soma, and its loss in testis. Despite extensive degeneration and down-regulation of the neo-Y chromosome, we find that the expression of neo-Y gametologs can compensate for reduced expression of the neo-X. Moreover, previously autosomal genes with testes-biased expression are disproportionally lost from the neo-X, thus becoming exclusively Y-linked, likely to resolve sexual antagonism. Interestingly, these now Y-exclusive genes are primarily expressed after meiosis, raising the possibility that postmeiotic germline regulation may be a major source of sexual antagonism driving the architecture of sex chromosomes. Co-amplified neo-sex genes are predominantly expressed during meiotic stages of spermatogenesis, consistent with their amplification being driven to compete for transmission to the next generation. In summary, our results shed light on the various molecular processes operating on sex chromosomes, and how they evolve.

## RESULTS

### Germline progression at single cell resolution

After processing and filtering, the replicated scRNA-seq experiment yielded 10,990 cells that fall into 12 clusters (**Figure 1D**, **Supplementary figure 1A**). We identified the cell types of eight clusters using testes marker genes (**Figure 1E, Supplementary figure 1B**). Four clusters show low but similar expression to the spermatid clusters but lack obvious expression profiles of key marker genes, thus we removed them from our downstream analyses (**Supplementary figure 1B**). This results in 8 cell types that span spermatogenesis with germline progression roughly following a linear trajectory in the UMAP projection (**Figure 1E**).

Consistent with previous reports in *D. melanogaster* testes [35,60], overall expression across all chromosomes increases through germline progression and peaks during meiotic exit at one of the late spermatocyte cell types (**Figure 1F**, **Supplementary figure 2**). During this stage, many genes (including cup and comet genes) are upregulated in preparation for sperm individualization and maturation [61,62]. The X chromosomes arms consistently show lower expression than the autosomes (except for Muller F, which used to be an X chromosome [52,63]) even in the somatic tissue where the X is expected to be fully dosage compensated. Lower expression from the X in male tissues compared to female tissues has been interpreted as an adaptation of the X to female-biased transmission (i.e. feminization of the X).

Nevertheless, across the stages, the ancestral X, Muller A, typically has the highest expression while the neo-X has the lowest expression, consistent with incomplete dosage compensation of the neo-X (see below). Consistent with the neo-Y degenerating and becoming heterochromatic, the neo-Y is lowly expressed compared to all other chromosomes. However, while expression peaks at all chromosomes at late spermatocytes (**Figure 1F**), genes from the neo-Y have the largest fold-increase between early and late spermatocyte stages, revealing that the neo-Y is disproportionally upregulated upon meiotic exit (**Supplementary figure 2**) [64].

### Reduced expression of X linked genes through meiosis consistent with loss of dosage compensation

*D. miranda’s* unique sex chromosome architecture presents an opportunity to test the status of X chromosome regulation in the germline and to understand its evolution (**Figure 2A**). If meiotic X-inactivation is an evolved trait in the male germline, the neo-X - which is at an intermediate stage of evolving X characteristics - should show expression patterns still reminiscent of its recent autosomal history. That is, genes on the neo-X will be expressed at a level that is intermediate of autosomal (uninactivated) and X (inactivated) levels during meiosis. In contrast, if reduced meiotic expression simply reflects loss of dosage compensation instead of targeted inactivation, opposite expression patterns are expected (**Figure 2A**). Since the neo-X is still evolving towards becoming fully dosage compensated, it should show lower expression compared to the other X arms prior to meiosis. However, through meiosis all three X chromosomes will lower to similar expression levels as dosage compensation is lost.

**Figure 2.**
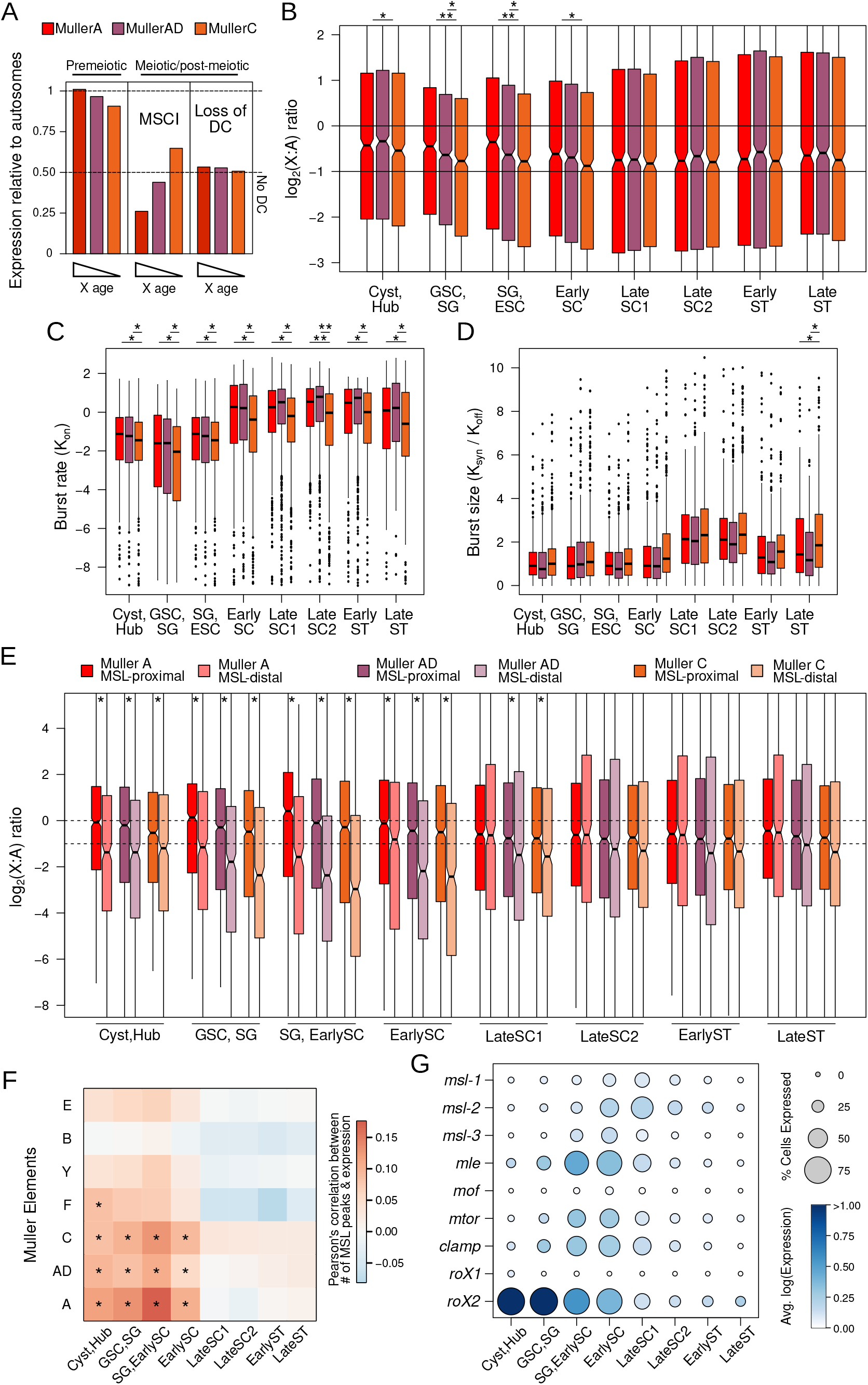
Gradual loss of dosage compensation through meiosis. **A.** Hypothetical models of MSCI versus loss of dosage compensation in Xs with different ages **B.** Distribution of X-to-autosome ratios across cell types. **C,D.** Estimated transcriptional burst rate (**C**) and burst size (**D**) for genes found on different X chromosomes. * = p < 0.001 and **= p < 0.000001, Wilcoxon’s Rank Sum Test. **E.** X-to-autosome ratios of genes based on their proximity to MSL-binding sites. Genes <2kb or >2kb away are classified as proximal or distal, respectively. **F.** Pairwise Pearson’s correlations between the number of nearby MSL binding sites and gene expression. * = p < 0.000001, Wilcoxon’s Rank Sum Test. **G.** Expression of the genes in or associated with the MSL complex across cell clusters. Size of the circle is proportional to the fraction of cells in the cluster with expression of the gene and the intensity of the color reflects the expression in log scale averaged across cells within the cluster.

To differentiate between these scenarios, we contrasted the expression of the three X’s relative to the autosomes throughout spermatogenesis (**Figure 2B**). We find that prior to meiosis (in GSC, SG, and SC), Muller C consistently has the lowest expression, with Muller AD showing intermediate expression levels between the two (**Figure 2B**). This stepwise difference in expression among X chromosomes during spermatogenesis is consistent with the gradual acquisition of dosage compensation over evolutionary time, where the ancestral X is fully dosage compensated, the neo-X shows partial dosage compensation, and Muller AD is in between (**Figure 2B**). After meiosis (in late SC and early ST), all three X chromosomes are becoming consistently lower expressed than autosomes (**Fig 2B**). In particular, expression levels of the older X chromosome arms begin to more closely resemble that of the neo-X across meiotic progression, equalizing at meiotic exist (late SC1). Most importantly, expression of all the X’s never drops below half that of the autosomes. This pattern supports the loss of dosage compensation as the mechanism underlying reduced X expression in testis and is inconsistent with the prediction that Muller C is evolving X-inactivation. Altogether, our results argue that the X’s are not inactivated or severely downregulated post-meiosis but are fully consistent with dosage compensation being lost during meiosis.

We further examined the transcriptional dynamics of chromosomes across spermatogenesis by inferring their expression kinetics [65,66]. Here, transcript production (K_SYN_) is modeled using two parameters, the frequency in which polymerases are engaged (burst rate, K_ON_) and shutdown (K_OFF_) [67]. Using this model, we find that burst rates indeed become elevated across meiosis for all X chromosomes. The neo-X has lower burst rate compared to the older X’s prior to meiosis consistent with incomplete dosage compensation. During and after meiosis, burst rates of the neo-X increase but remain consistently lower than that of the other X’s (**Figure 2C**). These patterns are, again, inconsistent with the model that the neo-X is in the process of evolving meiotic X-inactivation which predicts higher burst rates than the older X’s that would have extensive X-inactivation. Interestingly, the neo-X consistently produces more transcripts per burst (burst size), albeit not always significant; this leads to expression levels more similar to the two other X’s through meiosis despite relatively lower burst rate (**Figure 2D**). This elevated burst size of the neo-X may be a remnant of the neo-X’s recent autosomal history, as the autosomes also have elevated burst sizes (**Supplementary figure 3**).

### Loss of dosage compensation for genes proximal to MSL-bound regions across meiosis

To further evaluate the loss of dosage compensation across meiosis, we identified regions bound by the dosage compensation complex using ChIP-seq data [67,68] targeting the MSL3 protein (a component of the MSL complex). We identified 1513, 2762, 2115 MSL-bound regions for Muller A, AD, and C, respectively and then evaluated expression differences between genes proximal (<2kb) and distal (>2kb) to MSL-bound sites. Consistent with the MSL-complex inducing transcriptional upregulation of X-linked genes, in the somatic tissues and prior to meiosis, genes proximal to MSL-bound regions show significantly higher expression on all three X chromosomes (**Figure 2E**; Wilcoxon’s Rank Sum Tests p < 0.0001). However, as meiosis progresses and dosage compensation becomes lost, the expression of the two sets of genes become more similar and upon meiotic exit becomes statistically indistinguishable in most comparisons (**Figure 2A**; Wilcoxon’s Rank Sum Test p > 0.1). Thus, these patterns are again consistent with reduced expression of X-linked genes during spermatogenesis in Drosophila resulting from a loss of dosage compensation.

Transcriptional burst frequency also differs between X-linked genes proximal/distal from MSL-bound regions. Prior to meiosis genes near MSL-bound sites have significantly higher burst rate than those further away (**Supplementary Figure 4**; Wilcoxon’s Rank Sum Test p < 0.001) but the difference in burst rates decreases through and after meiosis. Moreover, we find significant positive correlations between expression levels of a gene and the number of MSL-bound sites nearby for all three X chromosomes prior to meiosis (Pearson’s correlation p < 10e-8); but through meiosis, such correlations are no longer observed (**Figure 2F**).

To specifically examine the activity of the dosage compensation complex, we looked at the expression of components of the complex and two accessory genes that are important for dosage compensation (*clamp* and *mtor,* **Figure 2G**). With the exception of *mof* and *roX1,* which have low expression across all cell stages (including the somatic cyst and hub cells and pre-meiotic stages), transcript levels of genes important for dosage compensation are intermediate to high in cell types that are dosage compensated. The long noncoding RNA, *roX2,* has the highest expression in the somatic cells and GSC, and gradually diminishes across germline progression. The others increase in expression peaking at the spermatocyte stages and decreases after. The expression pattern of these genes indicate that most genes involved in dosage compensation are active during early spermatogenesis and are gradually lost through meiosis, consistent with the loss of dosage compensation.

### Incomplete dosage compensation of the neo-X is exacerbated in the germline

Although our data show that all three X chromosomes of *D. miranda* are dosage compensated early in the male germline, the neo-X is consistently expressed at a lower level compared to the older X chromosomes, suggesting that dosage compensation on the neo-X may be incomplete. To specifically determine the extent to which the neo-X is dosage compensated in *D. miranda* males, we compared expression of orthologs between *D. miranda* and its sister species *D. pseudoobscura* where the neo-X, or Muller C, remains autosomal (**Figure 3A**). We used bulk RNA-seq data from whole testis and other sexed developmental stages and tissues including embryonic stages, larvae, head, and carcass from the two species. While genes on Muller A and Muller AD (X-linked chromosome arms in both species) show similar expression values in the two species, Muller C is under-expressed in several tissues of *D. miranda* males (**Figure 3B**, top and middle). Prior to zygotic genome activation, the embryo contains predominantly maternally deposited mRNA. Expectedly, in embryonic stage 2 (prior to zygotic genome activation in Drosophila), the difference between *D. pseudoobscura* and *D. miranda* expression of Muller C genes shows minimal bias with a median fold difference of 1 (**Figure 3B, bottom**). As zygotic expression ramps up through early embryonic development, Muller C genes become increasingly *D. pseudoobscura*-biased in expression, and stabilize in adult somatic tissues where the *D. miranda* alleles are expressed on average 0.769-fold less than the *D. pseudoobscura* alleles (**Figure 3B**). Therefore, neo-X linked genes are suboptimally dosage compensated across somatic cell lineages. In contrast, expression of Muller C genes in female tissues are highly similar between the two species, and the largest difference is a mere 0.1-fold reduction in *D. miranda* early embryos (**Supplementary Figure 5**).

**Figure 3.**
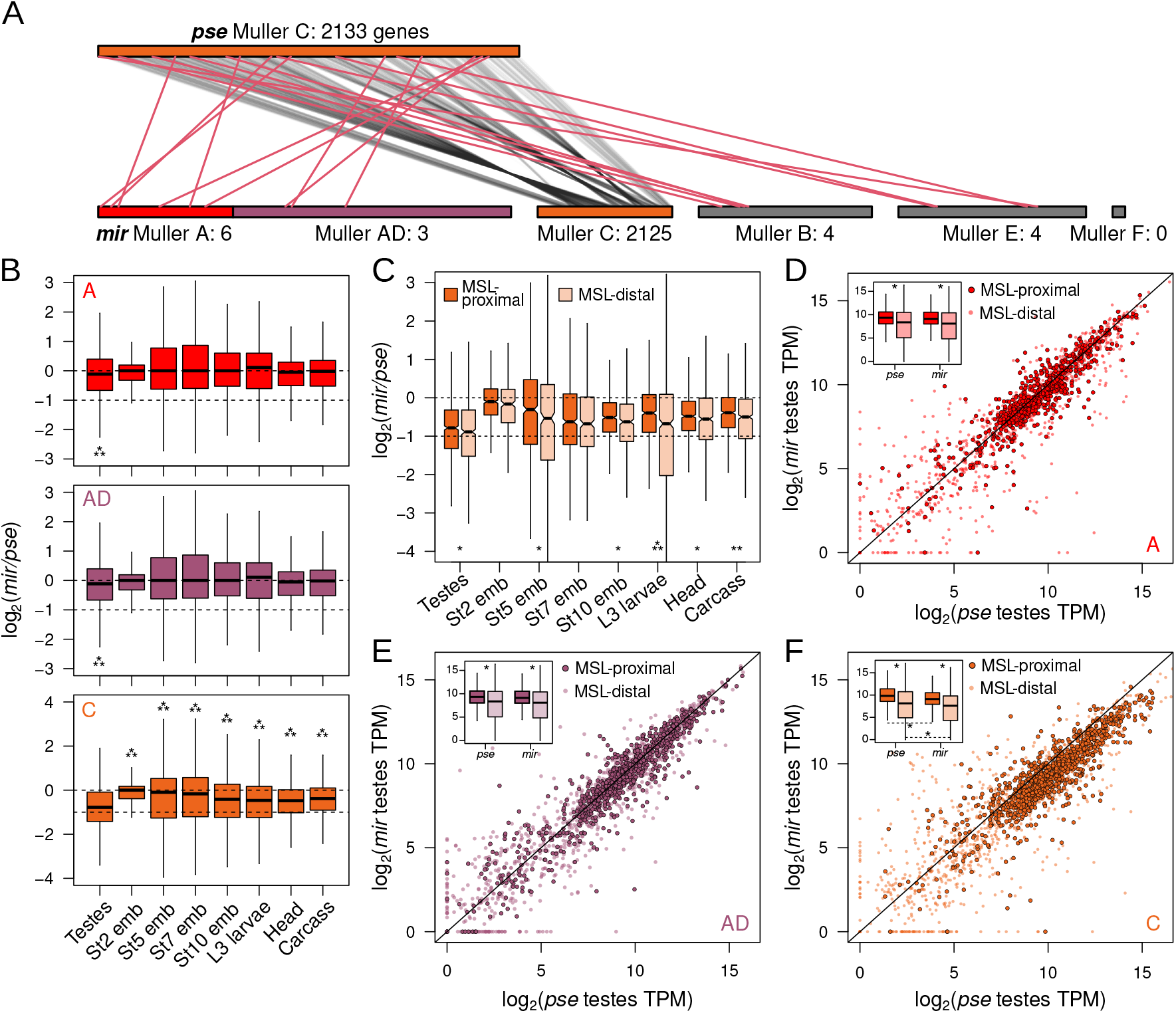
Divergence of X-linked expression. **A.** Gene movement and chromosomal positions of orthologs on Muller C in *D. pseudoobscura* before and after the formation of neo-X in *D. miranda.* Orthologous genes are connected by lines. Genes that migrated away (inter-chromosomal movement) from Muller C in *D. miranda* are in red. Number of genes from Muller C is noted next to the chromosome. Note, Muller C of D. pseudoobscura is drawn 4x the size for clarity. **B.** Log2(fold-difference) between *D. pseudoobscura* and *D. miranda* orthologs on different X-chromosomes and across different tissues and developmental stages. Note, the Y-axis differs on the bottom panel corresponding to Muller C. *** = p < 0.000001, Wilcoxon’s Rank Sum Test. C. Same as B, but Muller C genes are categorized based on their proximity to MSL-binding sites. * = p < 0.01, ** = p < 0.0001, and *** p < 0.000001. **D-F.** Correlation of orthologs expression on different X-chromosomes between *D. pseudoobscura* and *D. miranda.* Genes proximal to MSL-binding sites (<2kb) are displayed as darker circles with black borders. Diagonal line demarcates the identity line. The insets boxplots display the TPM distribution of MSL proximal and distal genes in the two species. * = p < 0.0000001.

As expected from the loss of dosage compensation through spermatogenesis (**Figs 1, 2**), testes show the most striking difference between the species. Here, *D. miranda* orthologs are expressed on average 0.565-fold of their *D. pseudoobscura* counterparts, a reduction that is a significantly greater than any other tissue (**Figure 3B** bottom, Wilcoxon Rank Sum Tests, p < 10e-8). However, while reduced expression of neo-X-linked orthologs is most extreme in the testes, the reduction is close to but nonetheless within 2-fold. Therefore, our results comparing the orthologs are consistent with the loss of dosage compensation across meiosis causing neo-X linked genes in some of the testes cell populations to have half of the expression of their autosomal counterparts in *D. pseudoobscura.*

Expression differences between orthologous genes proximal and distal to MSL-bound sites (in *D. miranda*) further confirm incomplete dosage compensation of the neo-X in *D. miranda* males, regardless of tissue type. Neo-X-linked genes distal to MSL-bound sites tend to be significantly more biased towards the *D. pseudoobscura* ortholog across most somatic tissues (**Figure 3C**; Wilcoxon’s Rank Sum Tests p < 0.01), consistent with reduced or a lack of dosage compensation. However, while proximity to MSL sites reduces the *D. pseudoobscura*-bias consistent with dosage compensation, MSL-proximal genes on the neo-X are still less expressed than their autosomal orthologs. The disparity between MSL-proximal and distal genes are most drastic in the L3 larvae, likely because the MSL-chip data were collected in the same developmental stage. However, even in L3 larvae, the MSL-proximal alleles are still on average 0.76-fold lower than their *D. pseudoobscura* orthologs. Therefore, even with wide-spread recruitment of MSL on the neo-X, dosage compensation is incomplete.

For the bulk testis samples, the difference between the MSL-proximal and -distal genes are marginal but significant with average fold differences of 0.580 and 0.540, respectively (Wilcoxon’s Rank Sum Test, p < 0.01). This slight difference is consistent with the loss of dosage compensation in many testis-derived cells (due to dosage compensation being abolished during meiosis). Looking across all three X chromosomes arms (**Figure 3C-E**), we find that genes near MSL-bound sites are enriched for highly expressed genes in both species. For Muller C (**Figure 3E**), we find that regardless of the proximity to MSL-binding sites, the *D. miranda* orthologs are consistently less expressed compared to *D. pseudoobscura* in the testes. Therefore, despite the presence of dosage compensation in some testes cell types, genes on the neo-X remain heavily under-expressed.

### Neo-Y gametologs mitigate neo-X dosage imbalance

Because of incomplete dosage compensation across tissues (and its loss during spermatogenesis), the neo-X of *D. miranda* appears to be in a state of suboptimal expression. This raises the question - how can males maintain fitness in lieu of this pervasive dosage imbalance? To address this question, we examined the neo-Y which, despite showing extensive degeneration of protein-coding genes, repeat accumulation and heterochromatinization, still harbors thousands of genes (gametologs) previously found on Muller C. Note that these gametologs are generally under reduced purifying selection and harbor an excess of deleterious amino acid mutations, and many contain premature stop codons or frame shift mutations. We identified gametologs on the neo-sex chromosomes and found that 1692 genes maintain both neo-Y and neo-X gametologs, 339 neo-X-linked genes with no neo-Y gametologs (neo-X-only), and only 36 neo-Y-linked genes with no neo-X gametologs (neo-Y-only) (**Figure 4A**). As expected for a heterogametic chromosome that is degenerating and not dosage compensated, for most of the genes that maintain both neo-Y and neo-X gametologs the neo-Y alleles are downregulated compared to the autosomal orthologs in *D. pseudoobscura* (Supplementary **Figure 6**) and their neo-X counterparts (**Figure 4B**).

**Figure 4.**
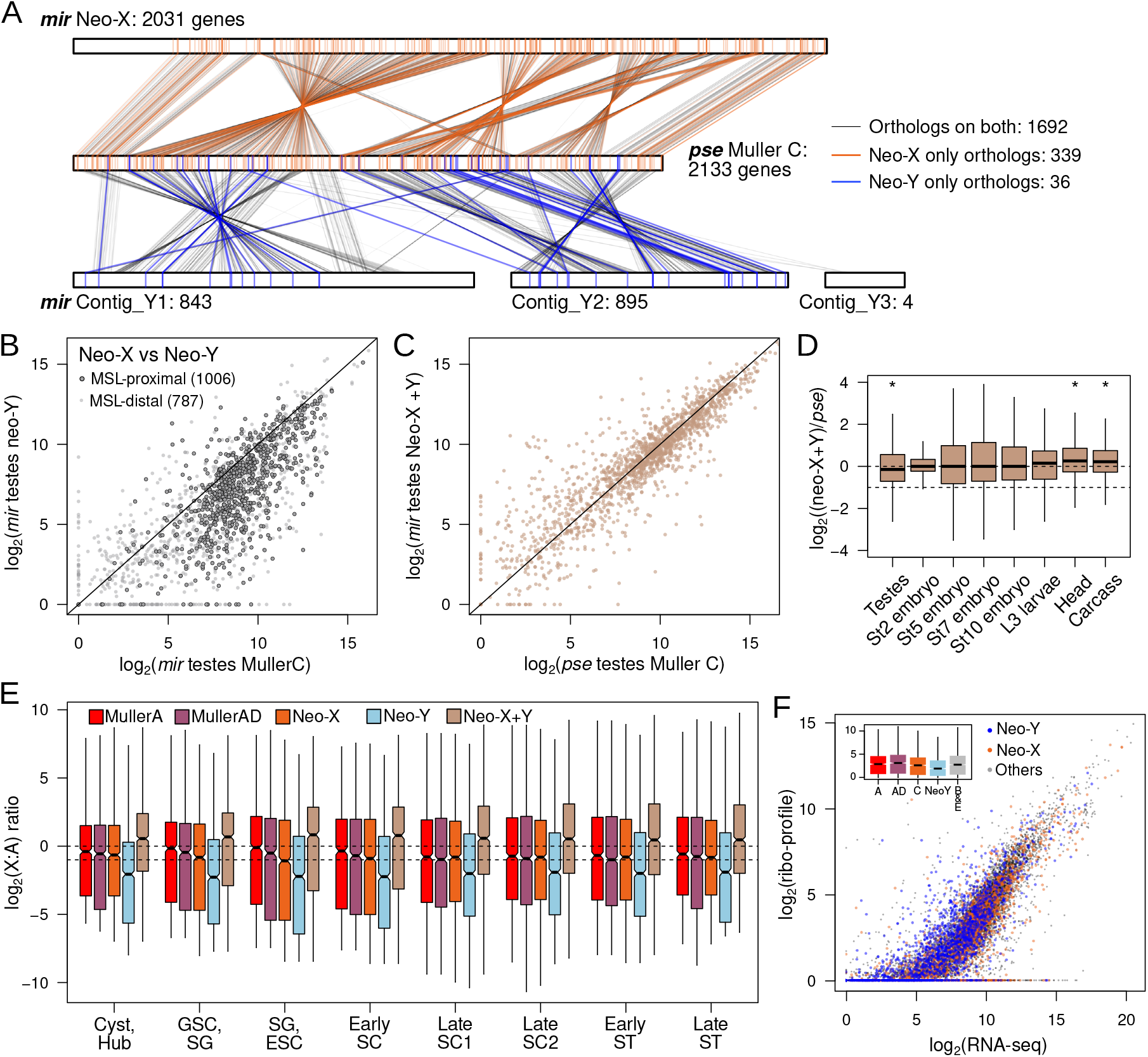
Y-linked gametologs mitigates expression and protein dosage imbalance. **A.** Synteny and positions of Muller C genes before (*D. pseudoobscura*) and after (*D. miranda*) the formation of neo-sex chromosomes. Muller C from *D. pseudoobscura* (middle) is sandwiched by the neo-X (top) and neo-Y (bottom) in *D. miranda.* Genes where both gametologs (n = 1692) are identified are connected with gray lines. Genes where only neo-X (n = 843) or neo-Y (n=36) gametologs are present are connected by orange or blue lines, respectively. Number of genes on each chromosome/contig is noted next to the chromosome labels. **B.** Testes expression correlation between neo-X and neo-Y gametologs. Diagonal line demarcates the identity line. **C.** Testes expression correlation between *D. pseudoobscura* gene on Muller C and the combined expression of neo-X and neo-Y gametologs in *D. miranda.* **D.** Log2(fold-difference) between the sum of *D. miranda* neo-X and neo-Y gametologs and D. pseudoobscrua Muller C-linked genes across different tissues and developmental stages. * = p < 0.000001, Wilcoxon’s Rank Sum Test. **E.** Sum of the neo-X and neo-Y transcript count across testes cell types in the scRNA-seq data. **F.** Correlation between the TPM from the RNA-seq data and ribosome profiling data from male larvae. The neo-X and neo-Y linked gametologs are plotted in orange and blue, respectively. Inset boxplot displays the distribution of the ribosome profiling reads across the chromosomes.

However, despite overall downregulation, many neo-Y gametologs are still highly expressed (**Figure 4B**) raising the possibility that neo-Y linked expression may mitigate dosage imbalance from incomplete dosage compensation, or its loss in the testes. Indeed, when we combined the expression of the neo-X and neo-Y gametologs, testes gene expression much more closely matches autosomal expression of genes on Muller C in *D. pseudoobscura,* resulting in a difference of only 0.92-fold (**Figure 4C and D**). In fact, this does not appear to be specific to testes, as dosage is generally restored to autosomal levels across all somatic tissues when expression from both the neo-X and neo-Y are combined (**Figure 4D**). This suggests that expression from the neo-Y linked gametologs ensures proper transcript dosage when dosage compensation is sub-optimal or not fully evolved.

However, expression of neo-Y alleles alone may not be sufficient to restore dosage at the protein level, since many Y-linked alleles have accumulated premature stop codons [68,69] and may not be translated. We therefore further evaluated translation of these transcripts using ribosome profiling (**Figure 4E**). Because premature stop codons lead to rapid degradation of mRNA via the nonsense mediated decay pathway [70,71], we expect nonfunctional transcripts from the Y to have low contribution to the translating mRNA. We indeed find that many of the neo-Y gametologs show high ribosome occupancy suggesting they are actively translated (**Figure 4E**). Unsurprisingly, the amounts of translating neo-Y transcripts are significantly lower than all other chromosomes (**Figure 4E, Supplementary figure 7A**), however they supplement corresponding neo-X gametologs, increasing the total translation rate of Muller C genes to near autosomal levels. Therefore, with incomplete dosage compensation of the neo-X, the neo-Y provides an additional source of genic and downstream protein production. Curiously, ribosome occupancy on the neo-Y gametologs appear to be uniquely elevated when normalized by transcript abundance (**Supplementary figure 7B**) and may suggest that the rate of translation is further enhanced by additional mechanisms.

### Neo-Y as an environment to resolve sexual conflict

Despite overall reduction in expression and gene loss, the neo-Y harbors 36 genes where the neo-X gametologs are lost. These neo-Y-only genes represent a set of genes that have become male-exclusive. Unlike the bulk of neo-Y linked genes which have lower expression compared to their neo-X-linked and autosomal counterparts (**Figure 4B and Supplementary Figure 6**), these neo-Y-only genes are overrepresented with genes highly expressed in the testes (**Figure 5A**; Wilcoxon’s Rank Sum Test p < 0.005). Interestingly, these genes are similarly highly expressed in *D. pseudoobscura* testes (where they are autosomal) and lowly expressed in females (**Figure 5B**), suggesting an ancestral function in testis. In addition to the neo-Y-only genes, we further identified 115 genes where the neo-Y gametolog is significantly higher expressed than the neo-X gametolog (neo-Y-biased). Similarly, we find that these neo-Y biased genes also show significant testes biased expression in *D. pseudoobscura* where they are still autosomal, although not to the same extent as the neo-Y only genes (**Figure 5B**). Reciprocally, strongly testes-biased genes in *D. pseudoobscura* are over-represented for neo-Y-only and neo-Y-biased genes in *D. miranda* (**Figure 5B**). These results reveal that the X is becoming demasculinized, losing testes genes to the Y chromosome.

**Figure 5.**
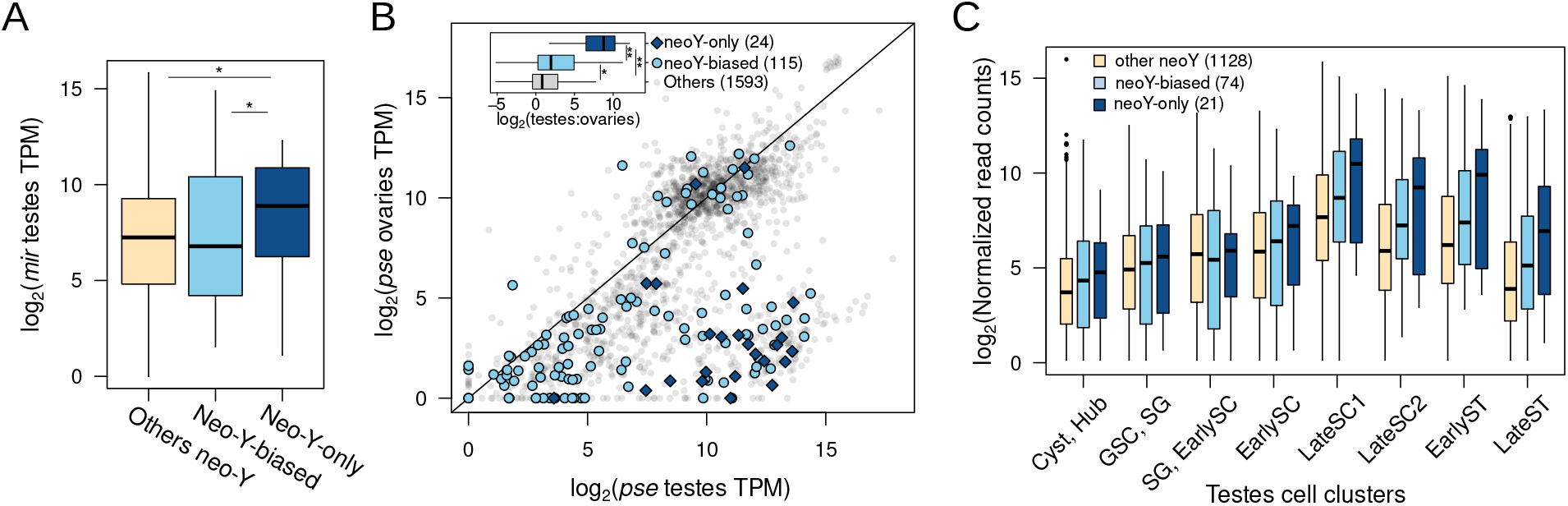
Y-biased and exclusive genes are enriched for testes-biased genes resolving sexual antagonism. **A.** Testes expression of neo-Y genes that show expression bias (neo-Y-biased) or have neo-X gametologs (neo-Y-only). * = p < 0.0001. **B.** Expression correlation of Muller C-linked genes between testes and ovaries in *D. pseudoobscura.* Genes where the neo-Y gametolog is more highly expressed than the neo-X gametolog in *D. miranda* testes (neoY-biased) or has only Y-linked gametologs (neoY-only) are labeled by light blue circles and dark blue diamonds, respectively. Inlet displays boxplots of the log2(fold-difference) between *D. pseudoobscura* testes and ovaries for the different gene categories. * = p < 0.001, ** = p < 0.00001, Wilcoxon’s Rank Sum Test. **C.** Expression of Y-linked genes across testes and spermatogenesis cell types showing elevated and lasting post-meiotic expression.

To specifically understand the germline regulation of these neo-Y-only and neo-Y-biased genes, we looked at their expression across the different testes cell clusters from the single cell data (**Figure 5C**). We find that compared to other Y-linked genes, neo-Y-only genes consistently show the highest expression across all cell types. Strikingly, their expression disproportionally increases through meiosis, peaking at late spermatocyte and remains high upon meiotic exit and sperm individualization. Therefore, late and postmeiosis appear to be the primary stages in which these neo-Y are most active.

After the formation of neo-sex chromosomes, the neo-Y is transmitted exclusively through males and therefore expected to become masculinized. While genes with important male function may be preferentially retained on the Y, it is unclear why they are lost or downregulated on the X. Genes with male-limited function could be lost on the X by random drift. Alternatively, these genes could represent sexually antagonistic alleles that are male beneficial and female detrimental and the neo-X linked copies are lost to eliminate deleterious effects in females. The formation of the neo-Y chromosome provides an opportune environment to resolve this conflict by allowing such genes to become male exclusive, alleviating deleterious effects in females.

Previous studies have revealed an excess of genes with testis-expression migrating off the X chromosome in Drosophila [72], and MSCI was invoked to explain this exodus. To determine whether a similar migration of X-linked testes genes to autosomes occurred on the neo-X of *D. miranda,* we identified orthologs between *D. miranda* and its sister species *D. pseudoobscura* where Muller C remains autosomal. Interestingly, we do not find an excess of Muller C genes to have moved to autosomes, albeit the number of gene movements is small (**Figure 4A**). Interestingly, we find that Muller A appears to be the most frequent destination for genes previously on the Muller C, which is inconsistent with the need to escape X inactivation. The unique opportunity to resolve sexual antagonism provided by the neo-Y offers an alternative path for genes with testes function to escape the X: After an autosome becomes sex-linked forming neo-sex chromosomes, male-beneficial female-deleterious genes are lost on the X to become Y-exclusive. However, this Y-linkage is only a temporary solution as gene corrosion and regulatory interference and epigenetic conflict from transposable elements will eventually drive these genes to either migrate off the Y or be lost entirely.

### Co-amplified neo-sex genes are predominantly expressed during meiotic stages of spermatogenesis

While Y evolution is characterized by massive gene loss and degeneration of the Y chromosome, we recently found that a subset of genes became highly amplified on both the neo-X and neo-Y chromosome after they became sex-linked [73]. Genes that co-amplified on the neo-sex chromosomes were highly enriched for functions associated with chromosome segregation, chromatin organization and RNAi, suggesting that their amplification was driven by X vs. Y antagonism for increased transmission. Consistent with co-amplified genes functioning to interfere with Mendelian segregation, we find that expression of both co-amplified X and Y genes drastically increases as they enter meiosis (peaking in late SC 1; **Figure 6A, B**). Co-amplified genes continue to be expressed post-meiosis when sperm chromatin is being remodeled, another process that has been found to be repeatedly targeted by meiotic drivers [25,74–76].

**Figure 6.**
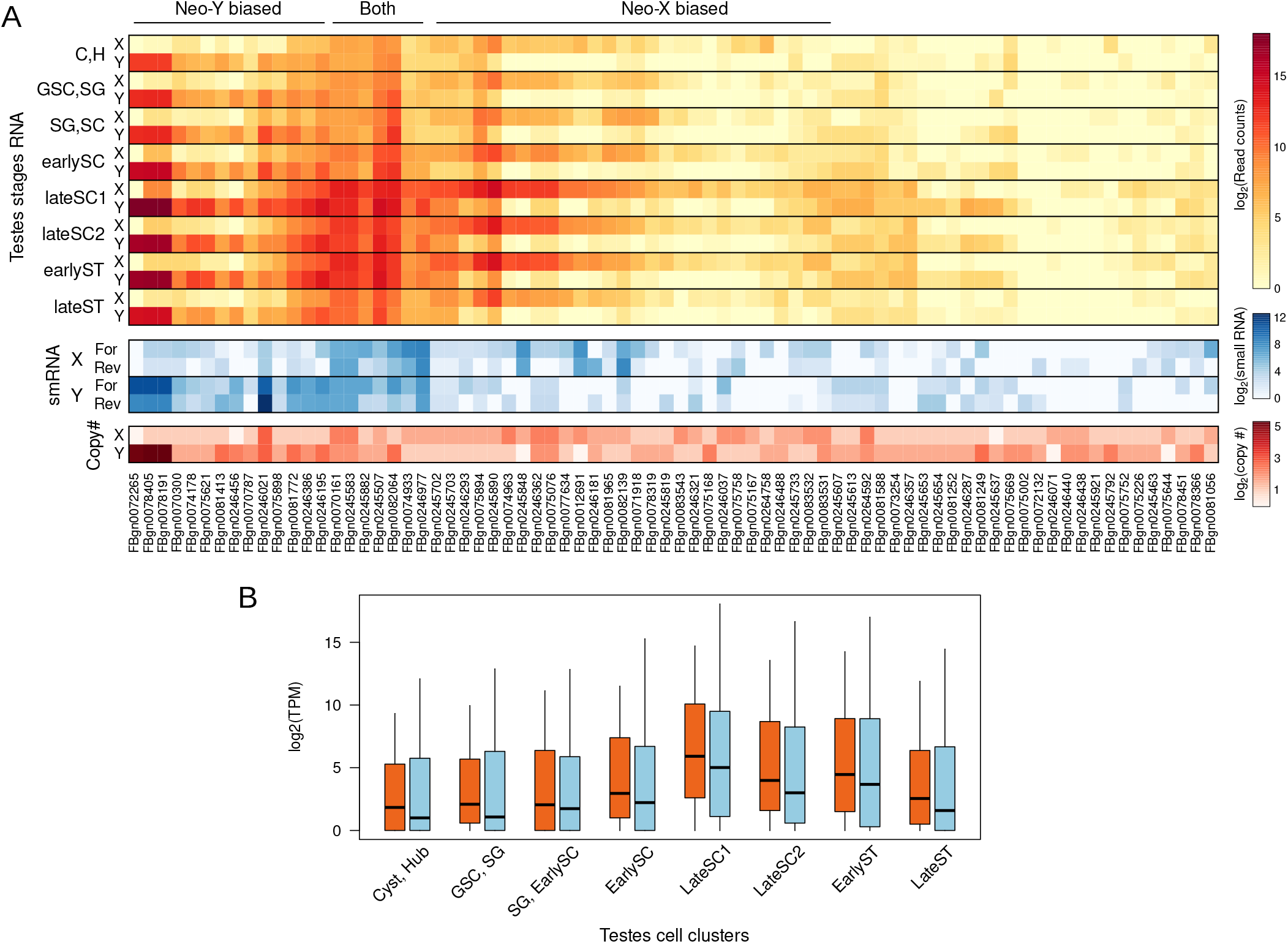
Expression of ampliconic genes on the neo-sex chromosome across germline progression. For each ampliconic gene family, the copy number, transcript abundance across cell types, and testes smRNA abundance are displayed in heat maps. For transcript abundance and smRNA, expression is summed across X- or Y-linked copies.

Expression patterns at co-amplified X/Y genes fall into four distinct clusters. Some genes are highly expressed from the neo-Y throughout spermatogenesis and peak during meiosis; they typically have very high copy number on the Y (up to 39 copies), and only few copies on the neo-X. These genes may have amplified on the neo-Y because of important testis-specific functions they perform during meiosis. The second cluster consists of genes that are highly expressed from both the neo-X and neo-Y chromosome. Their expression peaks during meiosis, and their copy numbers are more similar on the neo-X and neo-Y. We speculate that these genes are most likely to be involved in an ongoing, gene dosage-dependent conflict over segregation, where both X and Y copies are expressed during meiosis in a battle over transmission to the next generation. The third cluster consists of co-amplified genes predominately expressed from the neo-X copies; this could be ‘winning’ X-linked drivers pushing the population sex ratio towards females, as observed in wild *D. miranda* populations [73]. The last cluster contains co-amplified genes that show little expression from either chromosome. It is possible that these genes are now defunct relics of previous meiotic drive systems that are now effectively silenced.

Since the siRNAs pathway has been shown to suppress sex-linked meiotic drivers [77–79], we analyzed small RNA (smRNA) profile from bulk testes [73]. Interestingly, X-linked and Y-linked multicopy genes with high copy number and expression produce an abundance of both sense and antisense small RNAs (smRNA; **Figure 6**, **Supplementary Figure 8**). Further subdividing the smRNA reads based on their size, we find that the smRNAs are predominately distributed around 21bp indicative of siRNAs (and not simply RNA degradation products; **Supplementary Figure 9**). These results are consistent with the scenario that these sex-linked multicopy genes are reciprocally downregulating each other by producing siRNAs to bias their transmission into the next generation.

## Discussion

### Loss of dosage compensation through meiosis explains MSCI in flies

The status of meiotic sex chromosome regulation in Drosophila has been debated for decades. The first evidence for MSCI in flies, albeit indirect, was based on observations of reciprocal translocations between the X and autosome. Autosome-to-X translocations cause male sterility but not X-to-autosome translocations, suggesting that germline-essential genes translocated onto the X are downregulated [80]. In conjunction with claims of increased X-condensation in early spermatocytes, this led to the suggestion of MSCI in flies [80], but the cytological data were subsequently refuted [81]. More recent, direct evidence of MSCI includes reduced expression of transgenes inserted onto the X versus autosomes [29,33], and reduced expression of X-linked genes in carefully dissected testes tissues as assayed by qPCR and microarrays [31]. In conjunction with the observation that genes with testes-biased expression are under-represented on the X in flies, it has been proposed that MSCI drives the exodus of testes-biased genes off the X chromosome onto autosomes [30,34]. However, others have argued that the X is not globally inactivated [35], or that the reduced expression is not exclusive to meiotic stages but occurs even in pre-meiotic cells [36]. Discrepancies have been attributed to limitations in methodologies including limited number of genes quantified and tissue contamination in manual dissections, and single cell RNA-seq should be able to resolve the debate. Analyzing the meiotic cell types in adult Drosophila testes scRNA-seq, Witt et al found no evidence of MSCI [35]. However Mahadevaraju et al reported the presence of MSCI after analyzing scRNA-seq generated from developing testis discs in male larva [36]. Therefore, even with scRNA-seq the status of MSCI in *Drosophila* remains unclear.

We took advantage of *D. miranda’s* unique sex chromosome architecture to infer the presence of MSCI in Drosophila. *D. miranda’s* three X chromosome arms of different ages allowed us to determine whether the X chromosome is down-regulated, how this down-regulation occurs and how it evolves. Inconsistent with the X being inactivated during meiosis, X expression increases through meiosis. Both, transcript abundance (**Figure 1F**) and transcriptional burst rate (**Figure 2C**) of X-linked genes increases as germ cells progress through meiosis, although to a lesser extent compared to the autosomes. Furthermore, overall expression of the X’s never drops below half that of the autosomes. Prior to meiosis the neo-X consistently has the lowest expression of all three X’s, indicating incomplete dosage compensation. However, through meiosis expression of the older X’s simply drops to the same level as the neo-X and not below, as would be expected if MSCI is an evolved trait and the older X’s are more strongly downregulated (**Figure 2A**). These results collectively suggest that the X’s are losing dosage compensation during spermatogenesis, resulting in relatively lower expression compared to autosomes. This mechanism of relative X downregulation is fully consistent with both the findings in Witt et al [30,34] and Mahadevaraju et al [35,36]. Our results are also in partial agreement with the reports in Meiklejohn et al and Landeen et al that down regulation occurs across the testes and is not specific to meiosis [37], as the X’s in the somatic (Cyst and Hub) and early germline clusters (GSC and SG) all show expression levels that are significantly lower as would be expected for full dosage compensation (**Figure 2B**).

### Germline dosage compensation is less robust on the neo-X

Our analyses comparing X-linked genes proximal and distal to the MSL complex clearly show dosage compensation upregulating the former in many of the testis clusters until late meiotic cell types (late spermatocytes) (**Figure 1E**). While this is consistent with previous reports in *D. melanogaster* testes scRNA-seq dataset [80,81], the germline is thought to lack dosage compensation in flies. This is primarily based on the cytological observation that many of the MSL components are absent in the germline, and the male germline appears to lack X-chromosome specific H4K16ac [81,82]. However, we find that only *roX1* and *mof* are absent or lowly expressed in the germline tissues. *Rox1* and *rox2* have essential but redundant function in dosage compensation and *mof*, the histone acetyltransferase, appears to be lowly expressed even in other scRNA-seq datasets for somatic tissues that are expected to be dosage compensated [82,83], so low expression of *mof* may not necessarily reflect absence of activity. Thus, our data are consistent with MSL-dependent dosage compensation in the testis of *D. miranda,* but alternative means of achieving X upregulation in the GSC and spermatocytes without hyper-acetylation are possible. Three-strand DNA-RNA-hybrid structure (R-loops) were shown to be associated with hyper-transcription on the X in the absence of H4K16Ac [84]. Further, a recent comparison of the nuclear topology of germline stages reported that in the spermatogonia and spermatocytes, open chromatin on the X are more strongly associated and allow for higher expression than that of the autosomes. These studies both suggest mechanisms of upregulating the X in the absence of the full MSL complex.

Curiously, the (perhaps, non-canonical) dosage compensation in the germline appears to differentially upregulate the X arms. Even though the neo-X shows incomplete dosage compensation across all tissues (**Figure 3B**), even somatic ones, incomplete dosage compensation becomes much more severe in the early germline (**Figure 2B and E**), where the extent of dosage compensation tracks the age of the X’s. This suggests that germline dosage compensation is either weakest or being lost the quickest on the younger neo-X.

While dosage compensation of the neo-X appears most incomplete in testis, we also found that expression of the neo-Y alleles can increase overall expression from the neo-sex chromosomes (**Figure 4D**). By examining the expression of the gametologs on the neo-sex chromosomes in the scRNA-seq and comparing their expression to the ancestral autosomal expression in bulk RNA-seq data, we found that the neo-Y chromosome can mitigate suboptimal dosage compensation on the neo-X, regardless of the tissue types. Importantly, since neo-Y alleles are more highly expressed in testis compared to most autosomal tissues (**Supplementary Figure 10**), this means that overall expression from the neo-sex chromosomes in testis resembles that of the older X chromosomes more closely – i.e. only a moderate deficiency of X linked expression relative to autosomes in testis, similar to that of the older X chromosomes.

### Gene content and gene expression evolution resolves sexual and meiotic conflict of sex chromosomes

Gene content of sex chromosomes is non-random, and gene expression evolution and gene movement can contribute to sex chromosome specific expression patterns. We find that the neo-Y is becoming masculinized. Ancestrally testes-biased but autosomal genes have become either neo-Y biased in expression or neo-Y exclusive, indicating either downregulation or complete loss of the neo-X gametologs, respectively. Demasculinization of the X chromosome occurs, in part, due to the fact that the X chromosome spends twice as much time in females than males, and previous studies in Drosophila have revealed an excess of gene movement off the X chromosome of testis-expressed genes onto autosomes [6,42]. Here, we show that the neo-Y presents an intermediary for this movement as male-biased genes first transition to neo-Y biased or specific expression, alleviating the potential for these genes to be female-detrimental, thus resolving sexual conflict. However, as degeneration continues, these genes will inevitably be lost, further driving their movement onto autosomes. Interestingly, the formation of the neo-X in *D. miranda* is not associated with an excess of gene movements from Muller C to other autosomes (**Figure 3A** and **4A**); in fact, gene movement is small and Muller A appears to be the most frequent destination. Thus, autosomal migration likely occurs only after further degeneration of the neo-Y which provides a temporary but unstable solution to sexual conflict.

Sex chromosomes are also vulnerable to be involved in conflicts during meiosis. Meiotic drivers that try to cheat fair meiosis are prone to originate on sex chromosomes. Sex chromosome drive will skew the population sex ratio and select for suppressors on the other sex chromosome that are resistant to the distorter. If the distorter and suppressor are dosage sensitive, they would undergo iterated cycles of expansion, resulting in rapid co-amplification of driver and suppressor on the X and Y chromosome. We recently showed that meiosis and RNAi-related genes co-amplified on both the neo-X and neo-Y chromosome of *D. miranda.* Consistent with co-amplification of X/Y genes being driven to interfere with Mendelian segregation, we show that these genes are predominantly expressed during meiotic stages of spermatogenesis and postmeiotic stages, the cell types in which meiotic drivers are most likely to operate. Thus, sexual and meiotic conflicts may drive unique gene content and expression evolution on sex chromosomes.

## MATERIALS AND METHODS

### Testes disruption and single cell preparation

For each of the two replicates, five pairs of testes were dissected from *D. miranda* MSH22 in drops of cold PBS. Testes were transferred to a low retention Eppendorf tube containing a drop of lysis buffer (Trypsin L + 2mg/ml collagenase) followed by 30min of room temperature incubation with mild agitation. Then, the samples were first passed through a 50 um nylon mesh filter (Genesee Cat #: 57-106) with centrifugation at 1200 rpm for 7 minutes, and then twice through a 30 um nylon mesh (Genesee Cat #: 57-105). The resulting cell pellet was washed with 200ul of cold HBSS and pelleted again by centrifuging at 1200rpm for 7min. After removing the supernatant, the cells were resuspended in 20ul of HBSS. 5ul of the cell suspension were used for cell counting. 5ul were transferred to a slide for imaging with a Zeiss upright light microscope. Approximately 10000 cells were submitted for library preparation with the 10X Chromium platform at the Berkeley Vincent J. Coates Genomics Sequencing Lab followed by 100bp pair-end sequencing on Illumina NextSeq 2000.

### Single cell RNA-seq data processing

We used salmon (v1.5.2) [85] package to align the raw reads. We generated the index using a fasta file containing both *D. miranda* genic and transposable element transcripts. We then mapped the reads using the alevin program in salmon forcing 10000 cells with -- forceCells. Resulting files were then imported into R (v4.2.1) in Rstudio. We used Seurat (v4.0.5) [86] to process the read counts, cluster the UMIs, assign cell identities, generate UMAP projections, and aggregate cells within cell types. While there were initially 10990 UMIs, after clustering and cell type assignment using marker genes [35,36,60,62,87], we were unable to assign cell identity for 4 clusters that have low read counts (Supplementary figure 1). Cells (5101) in these clusters were removed, resulting in a final cell count of 5889. All subsequent analyses were done in Rstudio.

### Inferring transcription burst kinetics

We used the txburst software (https://github.com/sandberg-lab/txburst) and inferred K_SYN_, K_ON_ and K_OFF_ for genes in each cell cluster. In short, this software uses maximum likelihood estimation to estimate the three values based on the distribution of read counts for each gene across cells [65]. We only used genes with a TRUE value for the “Keep” column.

### MSL3-binding site and dosage compensation identification

The MSL-3 ChIP-seq is from ref [2]. We used bwa mem on default settings [88] (v 0.7.15) to align the chip and input reads. We then used macs2 [89] (v2.2.6) to call the enrichment peaks. We then used bedtools [90] (v2.26.0) closest to identified the gene closest to MSL3 peaks and vice versa. We used to bedtools intersect to identify genes overlapping or within 2kb of MSL3 peaks.

### Identifying orthologs and gametologs using reciprocal best hits

We downloaded the *D. pseudoobscura* (r3.2) CDS sequence from Flybase. For *D. miranda* we generated two sets of CDS files, one without neo-X genes and one without neo-Y genes. For each gene, the longest CDS is selected if there are alternative splice forms. We then used blastn (v2.6.0+) -task blastn to reciprocally blast the *D. pseudoobscura* CDS sequences with either sets of *D. miranda* CDS sequences. Best blast hit was defined as either having the highest e-value or longest blast alignment. Genes were identified as orthologs if they are reciprocal best hits of each other. The neo-Y gametologs are the Y-linked reciprocal best hits in the *D. miranda* CDS without the neo-X genes. The neo-X gametologs are the Muller C-linked reciprocal best hits in the *D. miranda* CDS without the neo-Y genes. Genes were further blasted to the *D. lowei* genome [91] to determine the ancestral chromosome location. Ampliconic genes were removed for orthology calls.

### Ribo-seq library preparation

We collected male third instar larvae and extracted total RNA using the RNAeasy Mini kit (Qiagen Catalog No. 74004). We digested the total RNA by micrococcal nuclease (Roche) with 3U/ug of total RNA. To collect the monosomes, we prepared 10-50% sucrose gradients in polysome gradient buffer (250mM NaCl, 15mM MgCl2, 20 U/ml Superase ln, 20ug/ml Emetine) with a GradientMaster (Biocomp Instruments). Monosome fractions were then collected after resolving and fractionating the gradient. We then extracted the RNA from the monosomes with a standard phenol/chloroform protocol and dephosphorylated the RNA by T4 polynucleotide kinase (New England Biolabs). We specifically excised RNAs spanning 28-34nt from the gel, and then performed two rounds of subtractive hybridization to remove ribosome RNAs as described. Small RNA libraries were then prepared following the standard Illumina protocol with TruSeq small RNA kit (Illumina Catalog No. RS-200). The ribo-seq libraries were sequenced at the Vincent J. Coates Genomics Sequencing Laboratory at University of California, Berkeley at HiSeq 2000 50bp SE.

### RNA-seq and ribo-seq data and processing

Embryonic and tissue-specific RNA-seq were from refs [92,93]. Reads were aligned using STAR [94](v2.7.10a) to their respective species genomes. The *D. miranda* genome is from ref [57] and the *D. pseudoobscura* genome (r3.2) is from Flybase. We then filtered the mapped reads for either the primary alignments or the unique alignments using samtools view –F 260 or –q 255, respectively. We then used featureCounts [95] (v2.0.3) in the Subread package to count reads over CDSs with featureCounts -p -M -t CDS. Because the X and Y can have large differential contribution to male and female transcriptomes, for normalization we used a modified version of Transcript Per Million with number of reads mapped to autosomal genes as the numerator: ((No. of reads mapped to a gene / CDS length*10e-6)) / (Sum of (No. of reads mapped to autosomal gene / CDS length) * 1000) * 10e-6).

## Notes

### Competing Interest Statement

The authors have declared no competing interest.

