## Supplementary Figure 1 for "Single cell RNA-seq in Drosophila testis reveals evolutionary trajectory of sex chromosome regulation"

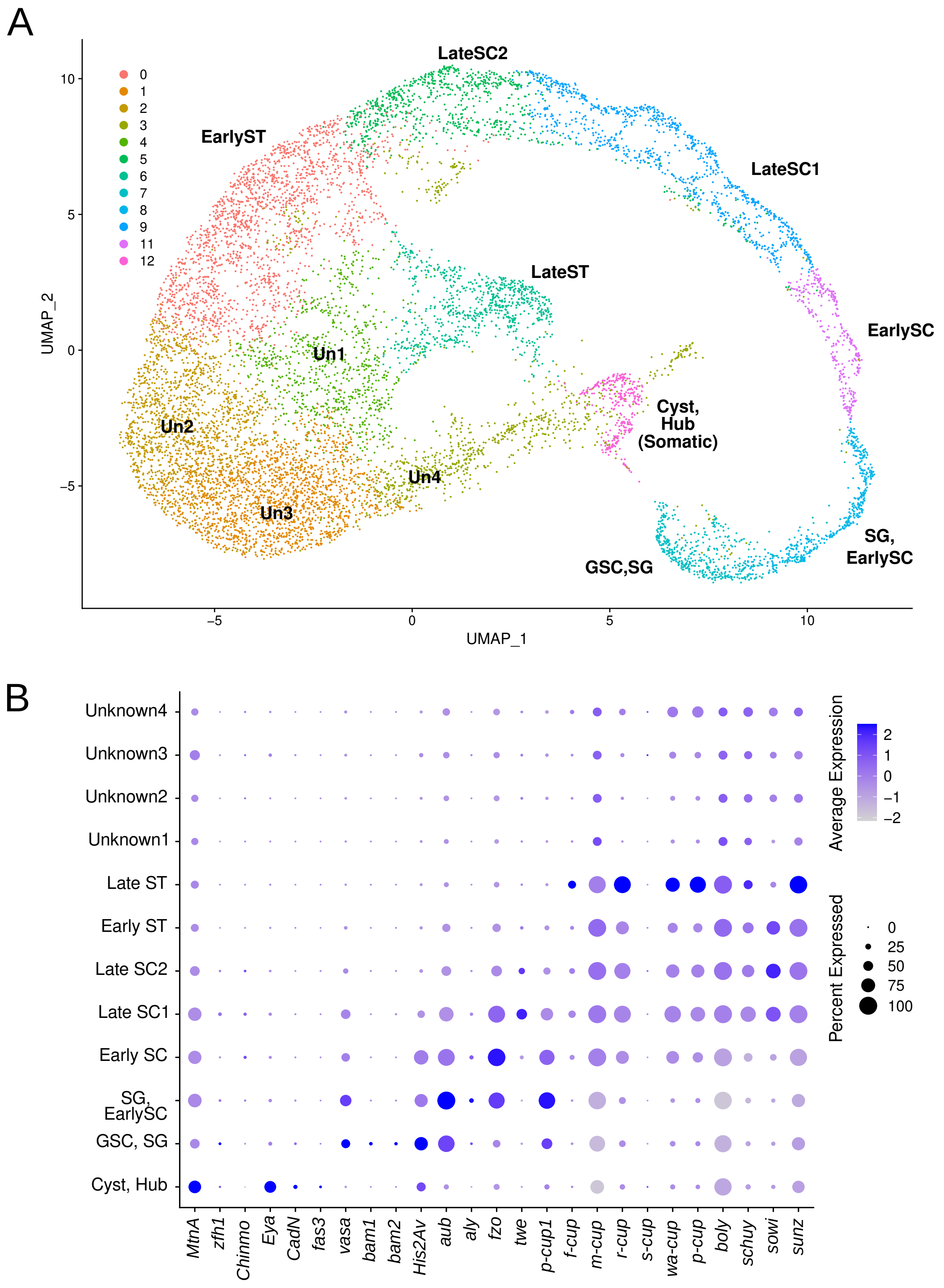


**Supplementary Figure 1:** A. UMAP projection of all UMI clusters. B. Expression of maker genes used to identify cell types across clusters.
