## Supplementary Figure 2 for "Single cell RNA-seq in Drosophila testis reveals evolutionary trajectory of sex chromosome regulation"

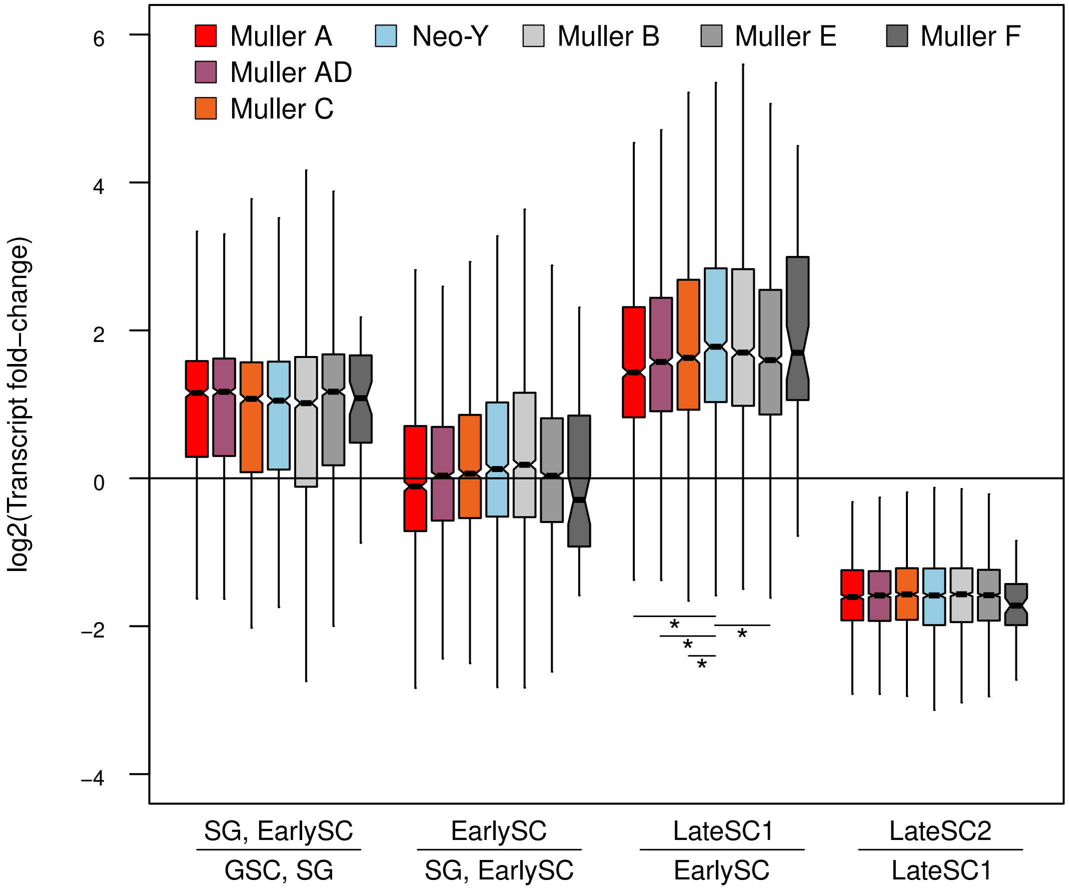


**Supplementary Figure 2:** Log scale distribution of expression fold differences between different cell stages for different chromosomes. * = p < 0.000001, Wilcoxon's rank sum test.
