## Supplementary figures and images for "Single cell RNA-seq in Drosophila testis reveals evolutionary trajectory of sex chromosome regulation"

### Supplementary Figure 3

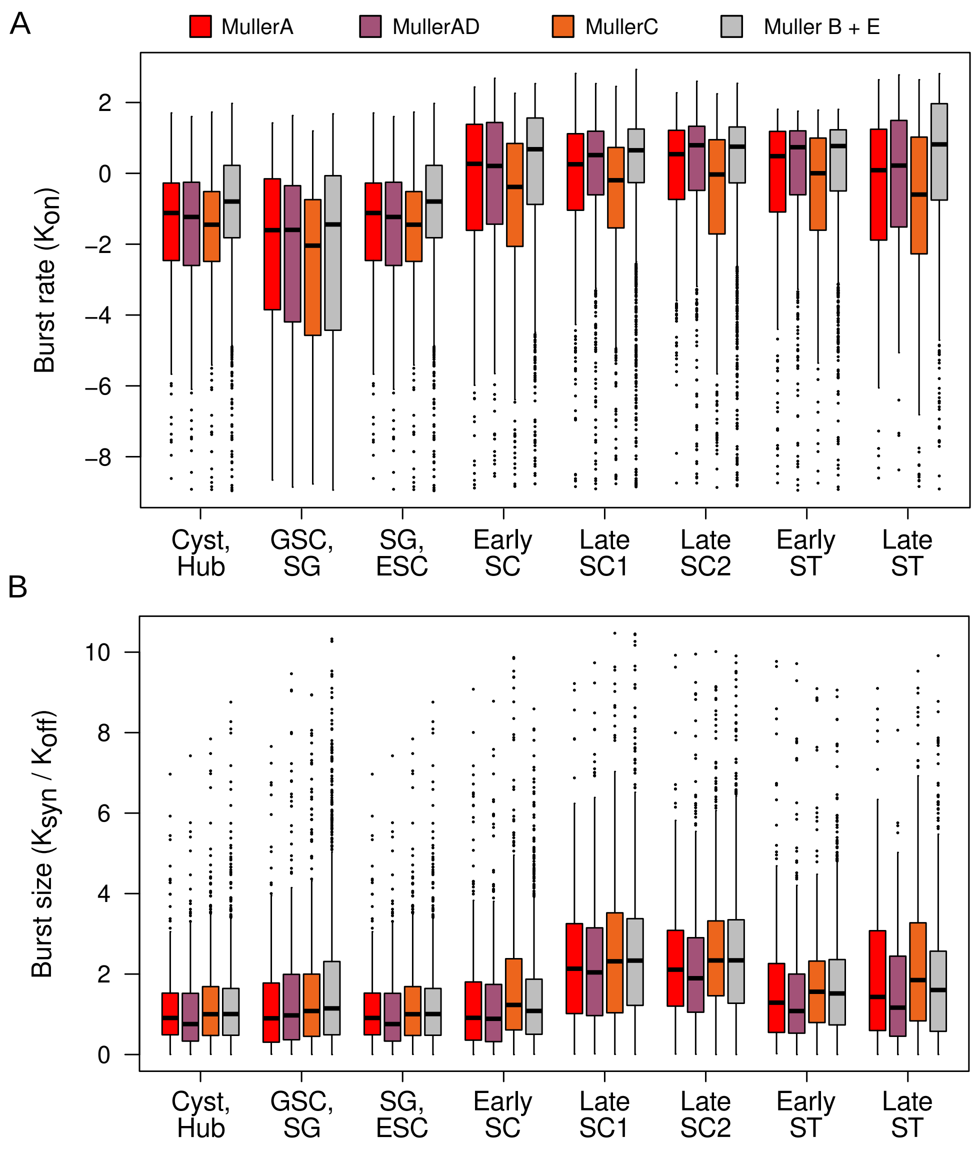


**Supplementary Figure 3:** Transcription kinetics relative to autosomes.

### Supplementary Figure 5

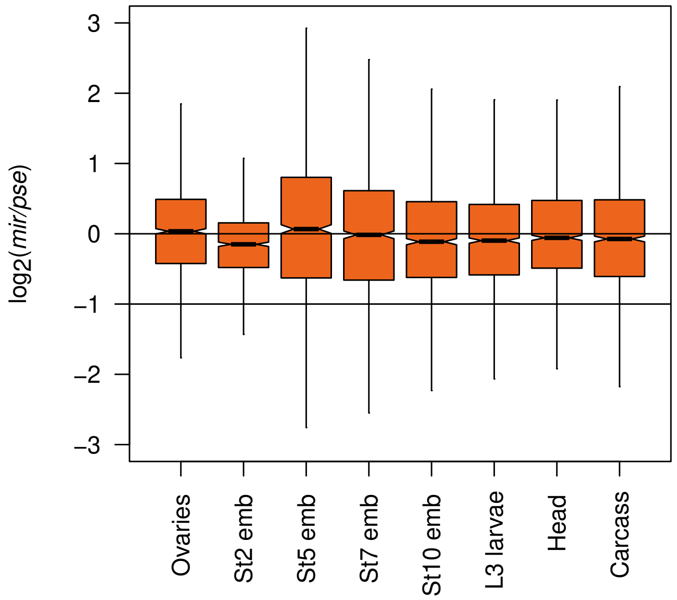


**Supplementary Figure 5:** Expression difference between Muller C orthologs in female tissues

### Supplementary Figure 6

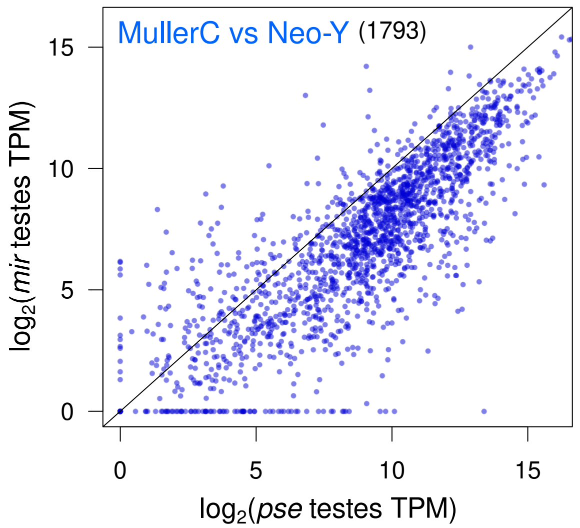


**Supplementary Figure 6:** Muller-C expression is D. pseudoobscura vs neo-Y expression in D. miranda.
