## Supplementary Figure 4 for "Single cell RNA-seq in Drosophila testis reveals evolutionary trajectory of sex chromosome regulation"

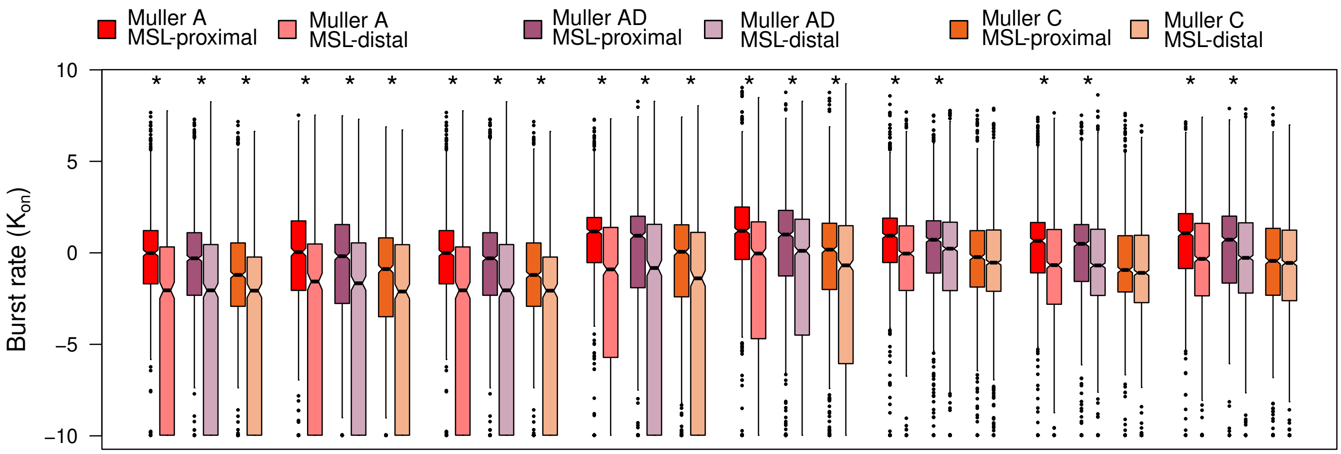


**Supplementary Figure 4:** Transcriptional burst rate of X-linked genes depending on the proximity to MSL-binding sites. * = p < 0.0001, Wilcoxon's rank sum test.
