## Supplementary Figure 7 for "Single cell RNA-seq in Drosophila testis reveals evolutionary trajectory of sex chromosome regulation"

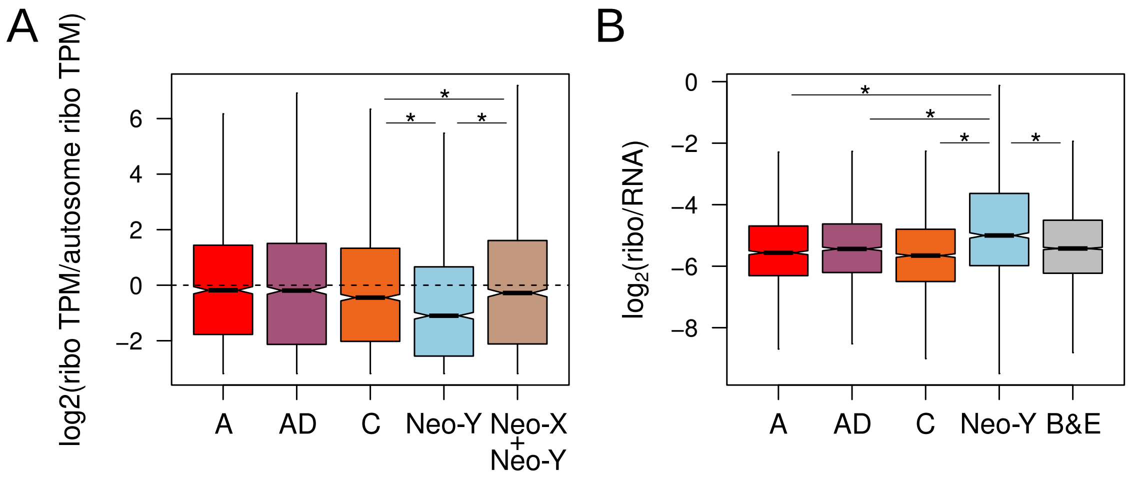


**Supplementary Figure 7:** Ribosome profiling and occupancy of *D. miranda* male larvae. Ratio of ribosome profiling reads of sex chromosomes to autosomes. Ribosome occupancy as measured by the ratio ribosome profiling reads over RNA-seq reads for the chromosomes. Elevated ribosome occupancy on the neo-Y gametologs could be a normalization artifact (higher rates of mis-mapping of shorter ribo-reads to the neo-Y), or suggest that the rate of translation is enhanced on the neo-Y by unknown mechanisms.
