## Supplementary Figure 8 for "Single cell RNA-seq in Drosophila testis reveals evolutionary trajectory of sex chromosome regulation"

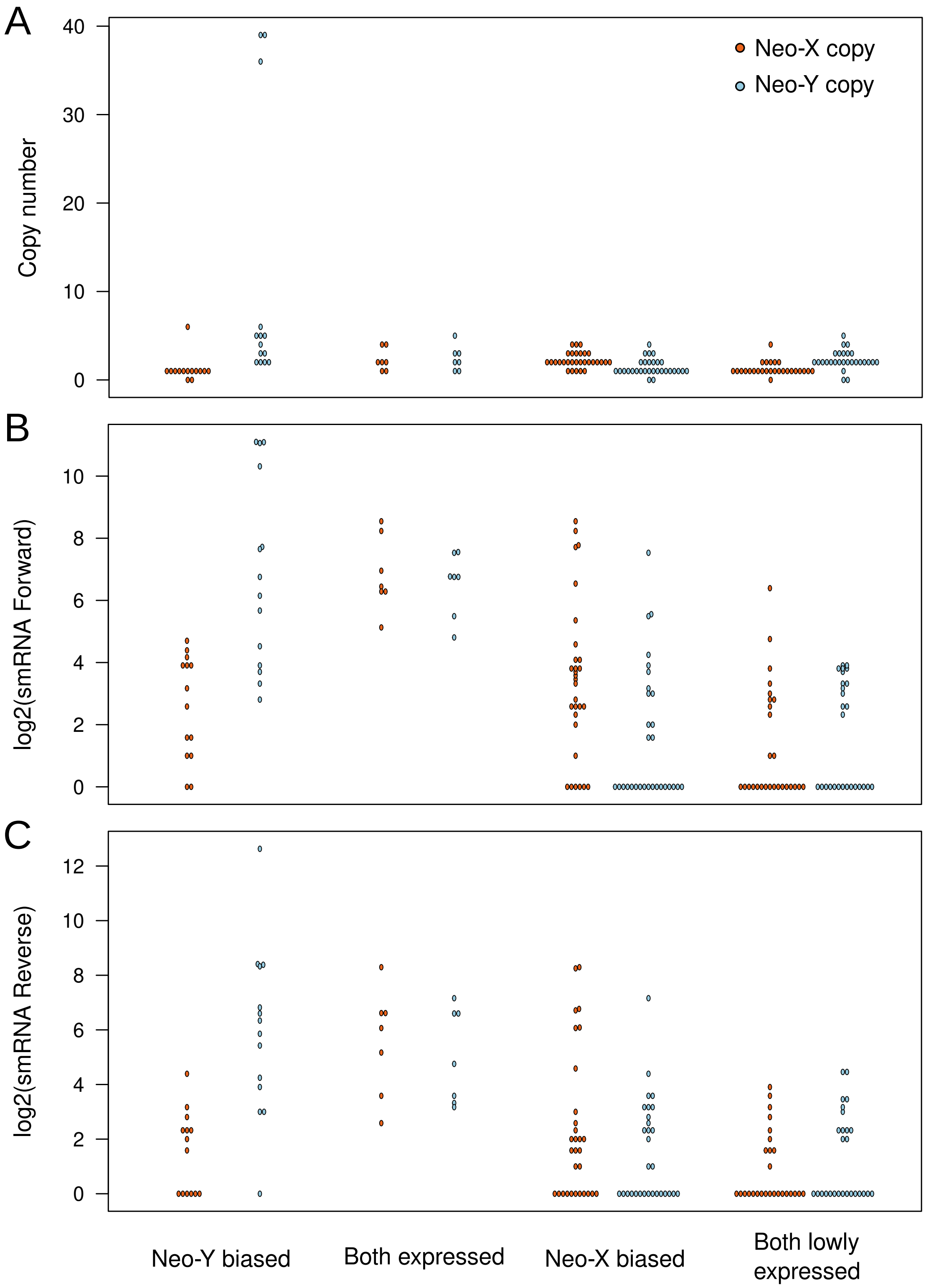


**Supplementary Figure 8:** Contrast between multicopy genes on the neo-X and neo-Y for copy number (A), abundance of sense smRNA (B), and abundance of antisense smRNA (C).
