## Supplementary Figure 9 for "Single cell RNA-seq in Drosophila testis reveals evolutionary trajectory of sex chromosome regulation"

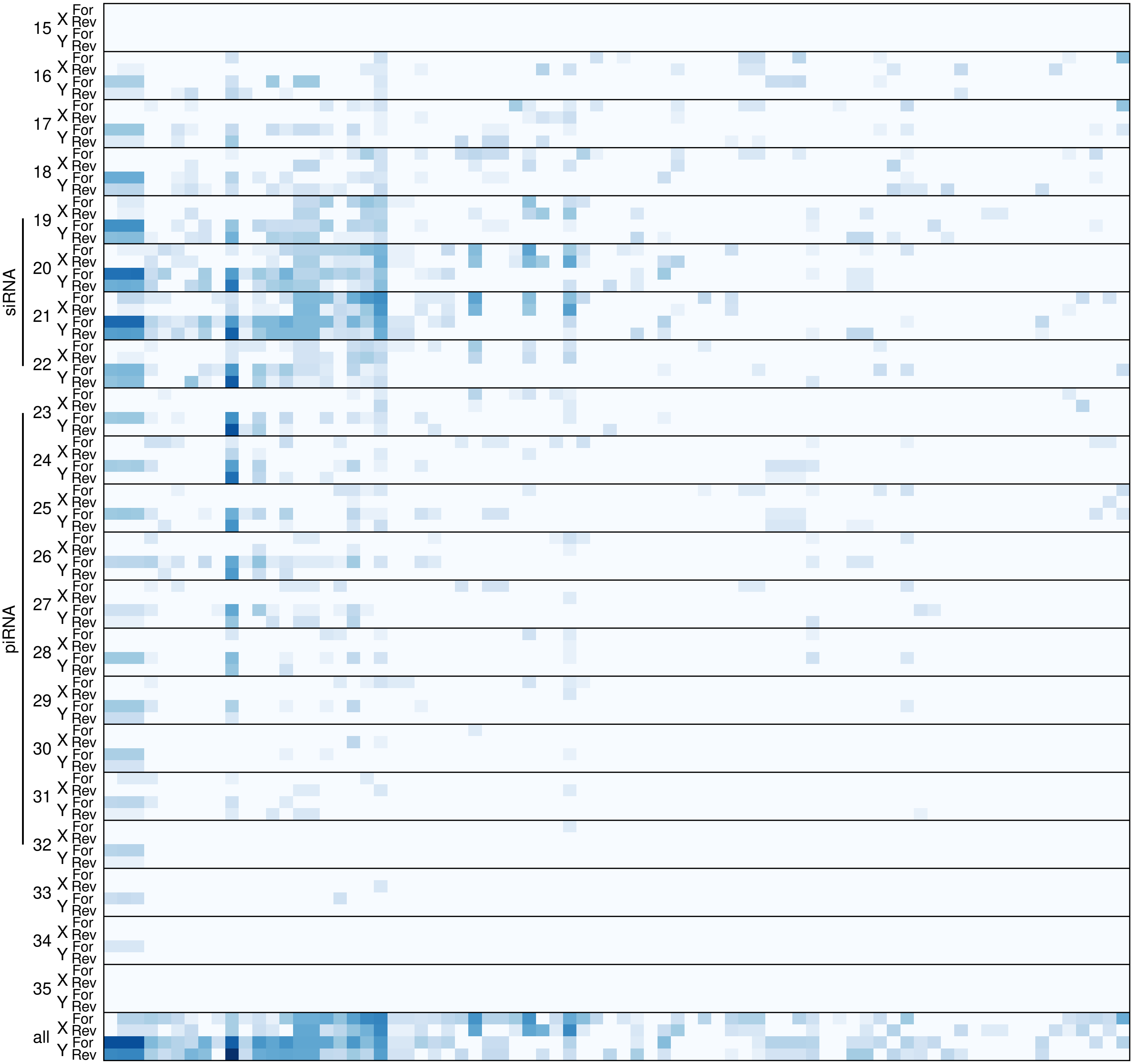


**Supplementary Figure 9:** smRNA abundance for X and Y linked multicopy genes broken down into fragment sizes corresponding to different types of small RNAs.
