## Supplementary Figure 10 for "Single cell RNA-seq in Drosophila testis reveals evolutionary trajectory of sex chromosome regulation"

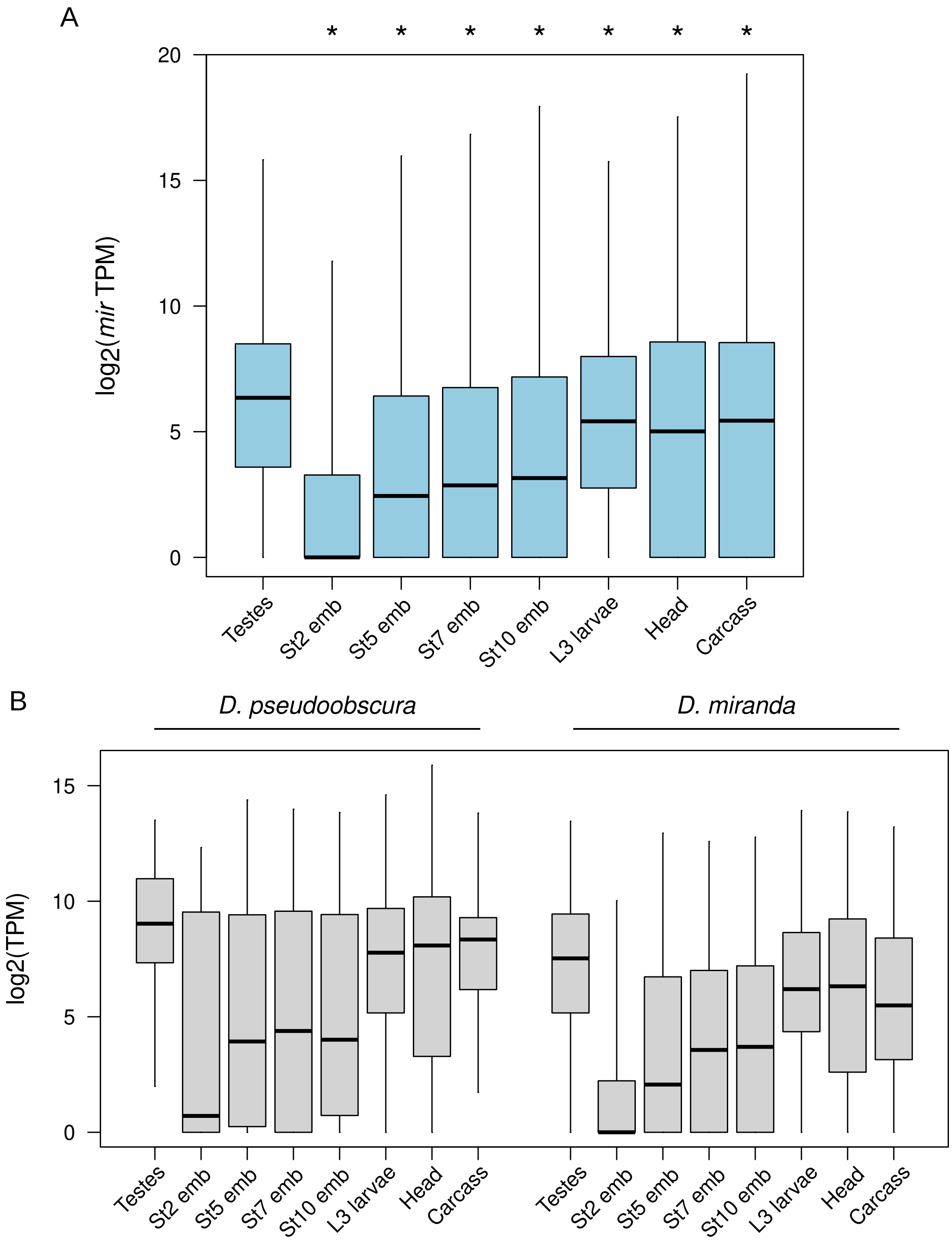


**Supplementary Figure 10:** A. Expression of neo-Y-linked genes across developmental stages and tissue types. * = p < 2.2e-16, Wilcoxon's rank sum test when compared to testes expression. B. Expression of Y-specific genes across developmental stages and tissue types in D. pseudoobscura and D. miranda.
